# Fine-tuning mechanical constraints uncouples patterning and gene expression in murine pseudo-embryos

**DOI:** 10.1101/2025.01.28.635012

**Authors:** Judith Pineau, Jerome Wong-Ng, Alexandre Mayran, Lucille Lopez-Delisle, Pierre Osteil, Armin Shoushtarizadeh, Denis Duboule, Samy Gobaa, Thomas Gregor

**Author notes:** Equal contribution, ordered alphabetically (Judith Pineau, Jerome Wong-Ng). Senior authors (Denis Duboule, Samy Gobaa, Thomas Gregor).

## Abstract

The interplay between mechanical forces and genetic programs is fundamental to embryonic development, yet how these factors independently or jointly influence morphogenesis and cell fate decisions remains poorly understood. Here, we fine-tune the mechanical environment of murine gastruloids, three-dimensional *in vitro* models of early embryogenesis, by embedding them in bioinert hydrogels with precisely tunable stiffness and timing of application. This approach reveals that external constraints can selectively influence transcriptional profiles, patterning, or morphology, depending on the level and timing of mechanical modulation. Gastruloids embedded in ultra-soft hydrogels (< 30 Pa) elongate robustly, preserving both anteroposterior patterning and transcriptional profiles. In contrast, embedding at higher stiffness disrupts polarization while leaving gene expression largely unaffected. Conversely, earlier embedding significantly impacts transcriptional profiles independently of polarization defects, highlighting the uncoupling of patterning and transcription. These findings suggest that distinct cellular states respond differently to external constraints. Live imaging and cell tracking imply that impaired cell motility underlies polarization defects, underscoring the role of mechanical forces in shaping morphogenesis independently of transcriptional changes. By allowing precise control over external mechanical boundaries, our approach provides a powerful platform to dissect how physical and biochemical factors interact to orchestrate early embryonic development.

## Introduction

Despite significant advances in developmental biology, the mechanisms that drive early embryogenesis—symmetry breaking, axis formation, germ layer specification, and tissue morphogenesis—remain incompletely understood. These processes arise from a complex interplay of genetic, biochemical, and mechanical signals (Collinet and Lecuit 2021), yet their precise interactions and temporal coordination remain elusive. Mammalian *in vivo* models are particularly challenging for such studies, as they are highly sensitive and constrained by the fact that these events occur post-implantation, making it difficult to disentangle the respective contributions of mechanical forces and biochemical cues. These limitations underscore the importance of controlled *in vitro* systems to systematically explore the role of physical and molecular factors in early development.

Recent advances in stem cell biology have enabled the creation of three-dimensional models known as gastruloids, which recapitulate key events of early mammalian embryogenesis. Gastruloids self-organize into aggregates that mimic symmetry breaking, anteroposterior (AP) axis elongation, and germ layer specification, providing a powerful platform for studying early developmental processes (Van Den Brink, Baillie-Johnson, et al. 2014; Turner, Girgin, et al. 2017; Beccari et al. 2018). Embedding these structures in extracellular matrix (ECM) substitutes, such as Matrigel, has demonstrated the critical role of mechanical properties in morphogenesis and cell fate determination (Veenvliet et al. 2020; Van Den Brink, Alemany, et al. 2020; Hamazaki et al. 2024; Muncie et al. 2020). However, the undefined chemical composition of Matrigel and its inherent mechanical properties are inextricably linked, making it impossible to separate these effects. Additionally, batch-to-batch variability poses significant challenges for quantitative studies (Hughes, Postovit, and Lajoie 2010; Vukicevic et al. 1992). Overcoming these limitations requires new approaches that combine controlled mechanical environments with high-resolution imaging to uncover how physical forces and biochemical signals coordinate development.

In this study, we leverage bioinert hydrogels with tunable stiffness to precisely control the mechanical environment of murine gastruloids. By systematically modulating both the stiffness and timing of embedding, we uncover how external constraints selectively influence transcriptional profiles, AP patterning, and morphology. Gastruloids embedded in ultra-soft hydrogels (*<* 30 Pa) elongate robustly while preserving both transcriptional profiles and AP patterning, mimicking the behavior of controls. In contrast, stiffer hydrogels (*>* 30 Pa) can disrupt polarization without altering gene expression, whereas earlier embedding significantly impacts transcriptional profiles independently of polarization defects. These findings reveal a surprising decoupling of transcriptional programs and AP patterning under specific mechanical conditions, challenging the conventional view of their tight coordination.

In addition to uncovering the uncoupling of transcriptional programs and patterning, our system minimizes sample movement during live imaging, enabling precise tracking of cell motility and morphogenesis. This advancement allowed us to identify impaired cell motility as a contributing factor underlying polarization defects in stiffer hydrogels. By providing precise control over mechanical constraints, our approach reveals how finely tuned environments can selectively influence distinct developmental outcomes. Together, these findings establish embedded gastruloids as a robust and versatile platform for probing the interplay between genetic, biochemical, and physical factors in early embryogenesis. By shaping our understanding of how mechanical environments guide developmental processes, this work offers novel insights into the regulatory principles of embryogenesis.

## Results

### Embedding in ultra-soft hydrogels allows reproducible gastruloid elongation

We developed an embedding procedure using a dextran-based hydrogel to investigate gastruloid elongation in a mechanically and chemically controlled environment (Figure S1A,E). By varying hydrogel concentrations from 0.7 mM to 1.5 mM, we achieved stiffnesses ranging from ∼1–300 Pa (Figure S1B-D), encompassing the range reported to support gastruloid elongation (Veenvliet et al. 2020; Van Den Brink, Alemany, et al. 2020). Importantly, this bioinert hydrogel minimizes extraneous signaling and variability, contrasting with traditional matrices such as Matrigel (Aisenbrey and Murphy 2020; Blache et al. 2022) (details in Methods and Figure S1).

We then compared the morphology of gastruloids embedded in hydrogels to those grown under standard culture conditions (Ctrl, no hydrogel). Gastruloids were embedded at 96 h post-seeding and analyzed at 120 h, the time frame during which the AP axis typically develops in our culture conditions.

Gastruloids embedded in hydrogels with concentrations below 1.0 mM successfully elongated, achieving approximately 80% of the medial axis length observed in controls (Figure 1A,B; Figure S2A-D). Interestingly, these embedded gastruloids exhibited a straighter morphology, as quantified by an increased straightness ratio, compared to controls (Figure 1A,C; Figure S2A-D). This suggests that while the mechanical constraints of the hydrogel did not prevent elongation, they counterbalanced bending forces, promoting straighter contours during elongation. Such straighter contours reduce shape variability, a key advantage for quantitative analyses and for interpreting morphological measurements.

**Figure 1:**
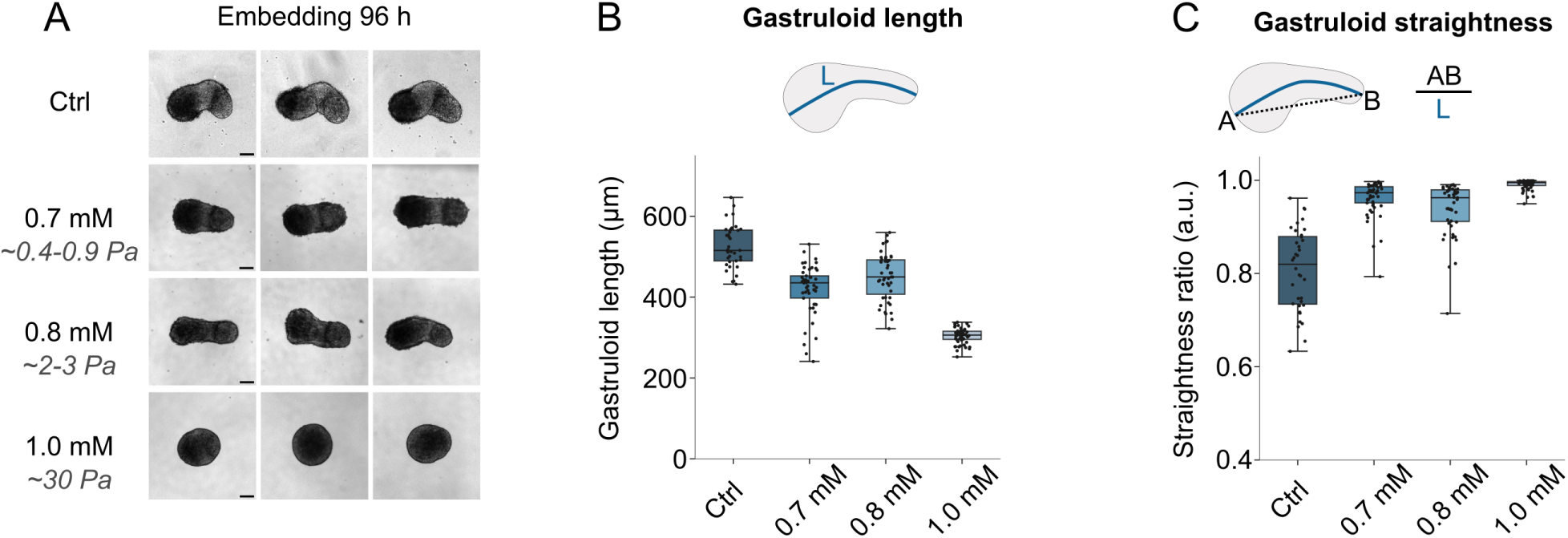
Impact of hydrogel stiffness on gastruloid elongation and morphology. (A) Representative bright-field images of gastruloids 120 hours after seeding. Gastruloids were grown in standard culture conditions (Ctrl, no hydrogel) or embedded in hydrogels with increasing concentrations (0.7 mM, 0.8 mM, or 1.0 mM) at 96 hours post-seeding. Scale bar: 100 μm. (B) Quantification of gastruloid medial axis length (*L*) at 120 h, showing moderately reduced elongation with increasing hydrogel stiffness. Data correspond to the conditions shown in (A). Sample sizes: Ctrl (*N* = 36), 0.7 mM (*N* = 50), 0.8 mM (*N* = 47), 1.0 mM (*N* = 56). (C) Straightness ratio of gastruloids from (B), revealing increased morphological straightness when hydrogel-embedded. Straightness is calculated as illustrated in the schematic.

In contrast, gastruloids embedded in higher stiffness hydrogels (1.0 mM) showed limited to no elongation (Figure 1B), and their straightness ratio approached 1 (Figure 1C), indicative of a lack of significant morphological changes. These findings demonstrate that the mechanical properties of the environment directly influence the elongation process, with ultrasoft hydrogels (*<* 1.0 mM) providing sufficient support for robust and reproducible elongation, while higher stiffness disrupts this process.

Notably, the embedding process also offers unique advantages for imaging and quantitative assays. By embedding gastruloids in a mechanically stable hydrogel, thermal fluctuations that often interfere with live imaging are minimized, enabling precise tracking of gastruloid dynamics. Additionally, the reduced variability in morphology, as indicated by straighter contours, facilitates reproducible quantitative measurements. Finally, hydrogel embedding provides a means to separate mechanical and chemical contributions to gastruloid development, highlighting its potential as an alternative to traditional assays employing Matrigel.

Together, these results establish a robust platform for studying gastruloid development in controlled mechanical environments, providing both physiological relevance and improved reproducibility for imaging and quantitative workflows.

### Mechanical embedding preserves gastruloid patterning and transcriptional profiles

During gastruloid development, the establishment of the AP axis is closely linked to axis elongation and patterned expression of key germ layer markers, such as BRACHYURY (BRA) and SOX2 (Blassberg et al. 2022; Turner, Hayward, et al. 2014; Van Den Brink, Baillie-Johnson, et al. 2014). These markers form a pole at the posterior end of gastruloids. To assess whether this patterning is maintained in hydrogels, we performed immunofluorescence staining and quantified intensity profiles along the AP axis (Merle et al. 2023).

Remarkably, a BRA/SOX2 pole was observed in all conditions, even in gastruloids grown in higher-stiffness hydrogels (1.0 mM), where elongation was impaired (Figure 2A, Figure S3A,C). To account for differences in fixation protocols between embedded and non-embedded gastruloids (see Methods), fluorescence intensities were normalized for both intensity and length, enabling a comparison of the spatial expression profiles. Gastruloids embedded in ultra-low stiffness gels (0.7-0.8 mM) exhibited AP expression profiles of SOX2 and BRA that closely resembled those of controls grown in standard culture conditions (Ctrl, no hydrogel) (Figure 2A-C, Figure S3A-D). However, gastruloids in 1.0 mM gels displayed spatial profiles that deviated from other conditions, likely due to weak elongation and challenges in consistently aligning the AP axis during imaging and analysis (Figure 2A-C, Figure S3A-D). These findings suggest that ultra-soft hydrogels preserve AP patterning comparable to control conditions.

**Figure 2:**
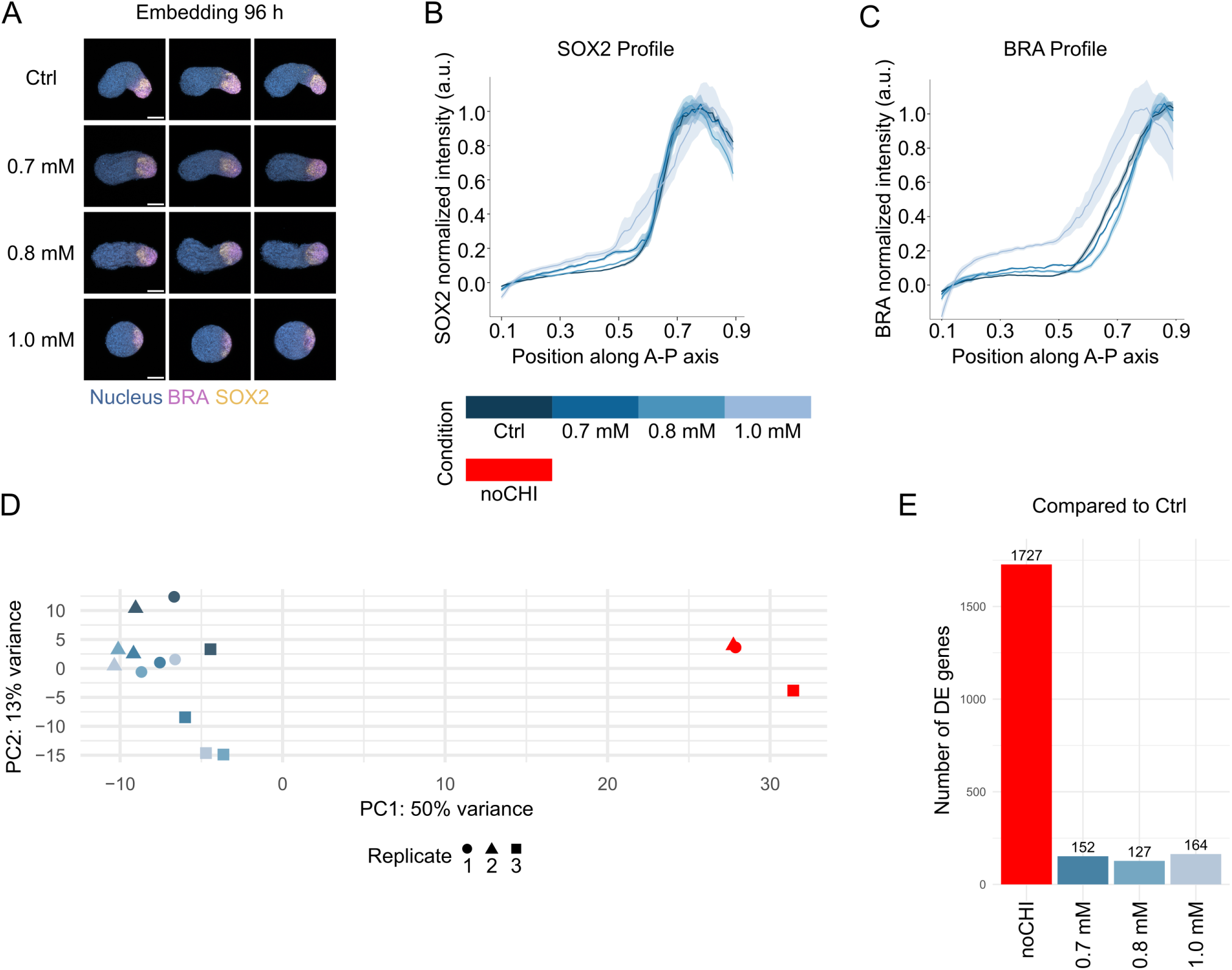
Gene expression patterning and transcriptional profiles in hydrogel-grown gastruloids. (A) Representative immunofluorescence images of gastruloids at 120 hours. Gastruloids were grown in standard culture conditions (Ctrl, no hydrogel) or embedded in hydrogels with increasing concentrations (0.7 mM, 0.8 mM, and 1.0 mM) at 96 hours post-seeding. Nuclei (blue), BRA (purple), and SOX2 (yellow) are labeled. Scale bar: 100 μm. (B, C) Normalized expression profiles (mean ± SEM) of SOX2 (B) and BRA (C) along the AP axis, demonstrating consistent expression patterns across conditions. Sample sizes: Ctrl (*N* = 18), 0.7 mM (*N* = 6), 0.8 mM (*N* = 19), 1.0 mM (*N* = 10). (D) Principal component analysis (PCA) of bulk RNA sequencing experiments (three replicates), revealing clear separation between Ctrl and noCHI (red), a negative control for gastruloid formation, while hydrogel-embedded conditions cluster near Ctrl. (E) Number of significantly differentially expressed (DE) genes compared to Ctrl, showing substantial transcriptional changes in noCHI and minimal changes in hydrogel-embedded conditions (0.7 mM, 0.8 mM, and 1.0 mM).

To evaluate whether gel embedding affects gene expression more broadly, we performed bulk RNA sequencing. As a negative control, we included gastruloids that did not receive a CHIR99021 pulse (noCHI), a condition under which gastruloids remain spherical and fail to establish germ layers (Van Den Brink, Baillie-Johnson, et al. 2014). Principal component analysis (PCA) on the top 500 most variable genes showed that 50% of the variance was explained by the difference between noCHI and all other conditions, while only 13% of the variance was attributed to differences among embedded and control gastruloids (Figure 2D, Figure S3E). Gastruloids embedded in ultra-soft hydrogels clustered closely with controls and no clear spatial separation was observed in the PCA, indicating high similarity between embedded and control conditions (Figure 2D).

The number of significantly differentially expressed (DE) genes between embedded and control gastruloids was small (Figure 2E, Supplementary table 1). Furthermore, we found no effect of gel concentration on transcriptional profiles (Figure 2D), suggesting that preventing elongation in higher-stiffness hydrogels (1.0 mM) does not strongly impact gene regulation. About half of DE genes are shared between all 3 gel concentrations (Figure S3F). This suggests that these differences may arise from the addition of the external boundary condition or the de-embedding process required for RNA sequencing sample preparation.

Altogether, these results demonstrate that embedding gastruloids in ultra-soft hydrogels preserves both AP patterning and transcriptional profiles, making this approach a viable alternative to traditional ECM-based matrices like Matrigel, which often exhibit batch-to-batch variability and ill-defined chemical compositions (Hughes, Postovit, and Lajoie 2010; Vukicevic et al. 1992). The minimal transcriptional and morphological deviations observed suggest that gastruloids grown in these conditions can be used interchangeably with those grown in standard culture, enabling the study of gastrulation in a mechanically controlled environment.

### Timing and stiffness reveal uncoupling of patterning and gene expression

As patterning and transcriptional profiles appeared robust to changes in the mechanical environment, we next sought to determine the limits of their establishment. Specifically, we tested whether increasing environmental stiffness to ∼ 300 Pa (Figure S1, S4) or applying mechanical constraints earlier in development could disrupt these processes.

To assess the impact of stiffness on patterning, we first examined whether gastruloids could organize a singular BRA/SOX2 pole at 120 h post-seeding when embedded in stiffer hydrogels at 96 h. Remarkably, most gastruloids formed BRA/SOX2 poles at 120 h, regardless of gel stiffness, with approximately 75% showing a singular pole even at higher stiffnesses (Figure 3A,B, S5). Notably, ∼ 20% of gastruloids had already established a pole by 96 h (Figure 3B, S5), suggesting that polarization begins either before or very shortly after embedding. These findings indicate that physical constraints applied after polarization is underway do not significantly disrupt this process.

**Figure 3:**
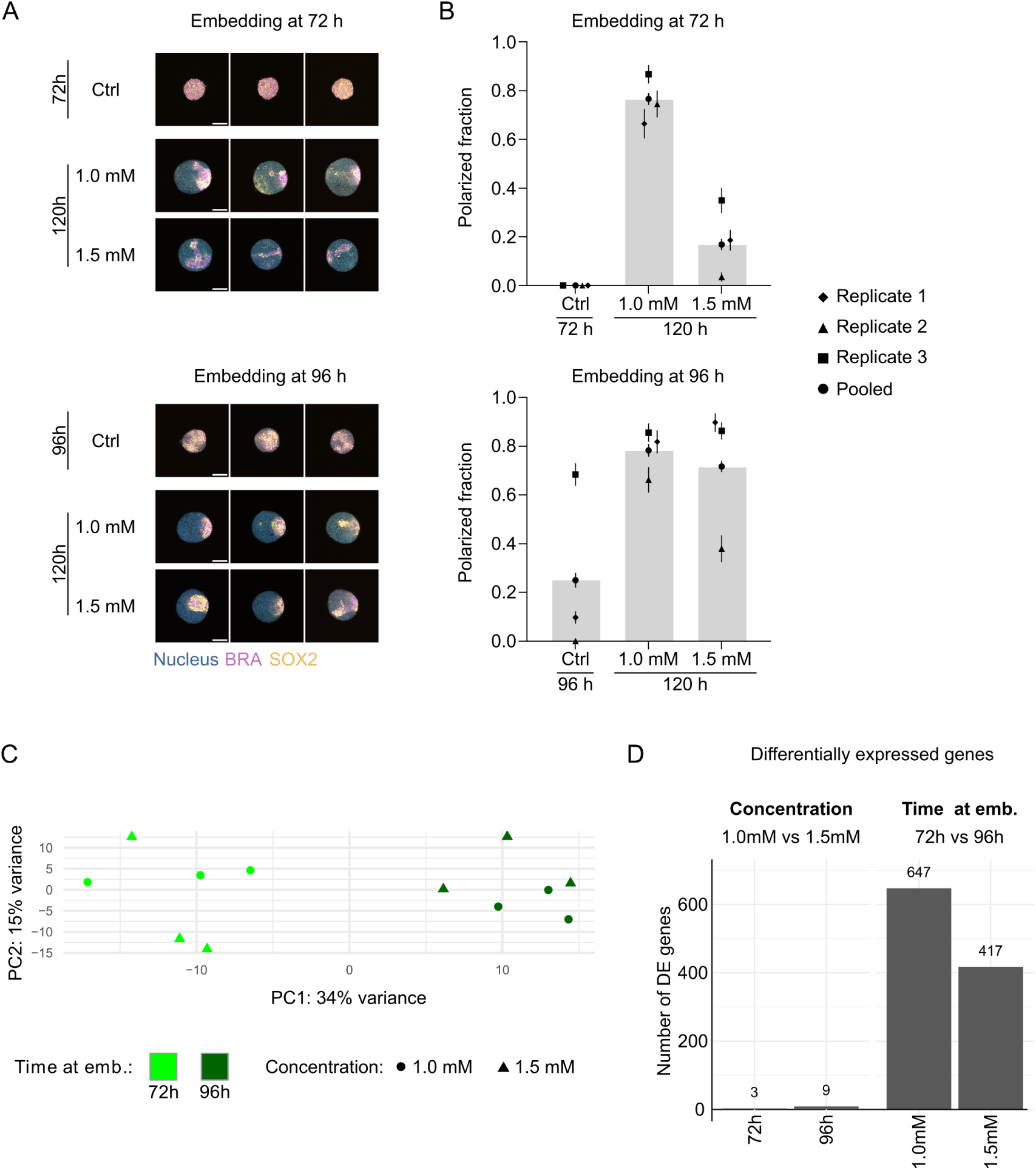
Uncoupling of patterning and transcriptional profiles. (A) Representative immunofluorescence images of gastruloids before embedding (72 h or 96 h) and at 120 h after seeding, following embedding in hydrogels at 72 h (top) or 96 h (bottom). Gel concentrations: 1.0 mM or 1.5 mM. Nuclei (blue), BRA (purple), and SOX2 (orange) are labeled. Scale bar: 100 μm. (B) Quantification of the proportion of gastruloids forming a unique BRA/SOX2 pole at 120 h or at the time of embedding. Data correspond to the conditions shown in (A) and include embedding at 72 h (top) or 96 h (bottom). Error bars were obtained from bootstrapping (see Methods). Replicates are shown in Figure S5. Sample sizes for embedding at 72 h: 72 h Ctrl (*N* = 20 for replicate 1; 18 for replicate 2; 22 for replicate 3); 120 h 1.0 mM (*N* = 15; 19; 22); 120 h 1.5 mM (*N* = 21; 27; 17). Sample sizes for embedding at 96 h: 96 h Ctrl (*N* =22; 23; 22); 120 h 1.0 mM (*N* = 21; 21; 22); 120 h 1.5 mM (*N* = 19; 21; 23). (C) Principal component analysis (PCA) of bulk RNA sequencing experiments, revealing transcriptional differences between gastruloids embedded at 72 h or 96 h in hydrogels of 1.0 mM or 1.5 mM concentrations. (D) Number of differentially expressed (DE) genes from bulk RNA sequencing, demonstrating minimal transcriptional changes between gel concentrations (1.0 mM vs 1.5 mM) at either embedding time and showing greater differences when comparing embedding times (72 h vs 96 h) within each gel condition.

Next, we investigated whether earlier mechanical constraints could impair polarization by embedding gastruloids at 72 h, a time point when no pole is established yet (Figure 3A, S5). When embedded in 1.0 mM gels (corresponding to 30 Pa), ∼ 80% of the gastruloids successfully formed a BRA/SOX2 pole by 120 h. However, this fraction dropped to ∼ 20% in 1.5 mM gels, suggesting that stiffer gels impose sufficient mechanical constraints to impair polarization establishment (Figure 3B, S5). Interestingly, gastruloids in 1.5 mM gels were ∼ 10% smaller on average (Figure S4), potentially indicating increased cellular compression or decreased cell proliferation. These results demonstrate that polarization establishment is sensitive to mechanical constraints during early developmental stages but remains robust once initiated.

To evaluate whether global gene expression was affected by these experimental conditions, we performed bulk RNA sequencing. Principal component analysis (PCA) revealed that samples clustered primarily by the time of embedding rather than gel stiffness (Figure 3C, S6). The first component (PC1) that separates samples by embedding time explained ∼ 34% of the observed variance in gene expression, whereas gel stiffness had minimal impact. For a fixed embedding time, only a small number of genes were differentially expressed between 1.0 mM and 1.5 mM gels (3 genes at 72 h, 9 genes at 96 h). In contrast, embedding at 72 h versus 96 h resulted in hundreds of differentially expressed genes, regardless of gel stiffness (647 genes for 1.0 mM, 417 genes for 1.5 mM). These genes included key regulators of embryonic development and stem cell differentiation, such as *Dppa5a*, *T/Bra*, and *Hoxa3* (Figure 3D, S6, Supplementary table 2).

Surprisingly, the inability of gastruloids embedded at 72 h in 1.5 mM gels to form a BRA/SOX2 pole did not correspond to significant transcriptional changes, as their global gene expression profiles were similar to those of gastruloids that successfully formed poles in 1.0 mM gels. Furthermore, embedding at 72 h induced major transcriptional changes irrespective of whether polarization was maintained (1.0 mM) or disrupted (1.5 mM). This unexpected result contrasts with the assumption that morphological polarization and transcriptional programs are tightly coupled, revealing instead that mechanical constraints can disrupt one without significantly affecting the other. These findings reveal that patterning and transcription can vary independently, providing strong evidence for their uncoupling under specific mechanical conditions.

Together, these results demonstrate that while polarization and transcriptional profiles are generally robust to mechanical constraints, their establishment can be selectively disrupted by the timing and stiffness of embedding. This uncoupling of polarization and gene expression highlights the distinct regulatory mechanisms underlying gastruloid patterning and transcriptional programs.

### Impaired cell motility in dense gel confinement

The minimal differences in transcriptional profiles between gastruloids embedded at 72 h in 1.0 mM and 1.5 mM gels (Figure 3C,D) contrasted sharply with their differing abilities to form a BRA/SOX2 pole (Figure 3A,B). Recall that 1.5 mM corresponds to a stiffness of approximately 300 Pa, and that there is an order of magnitude in difference in stiffness between 1.0 mM and 1.5 mM gels (Figure S1). This discrepancy suggested that defects in BRA/SOX2 pole establishment could arise from differences in cell motility, likely due to the increased mechanical constraints build-up in stiffer gels.

Long-term imaging and tracking of freely floating gastruloids is often hindered by translational and rotational movement during development. Embedding gastruloids in hydrogels, however, significantly reduces movement, even in ultra-soft hydrogels that support elongation (Figure 4A). This stabilization enables high-resolution imaging without requiring extensive image registration, addressing a major limitation of current imaging workflows. Existing solutions, such as micro-wells or holders, often impose size constraints or require labor-intensive post-processing steps (Beghin et al. 2022; Samal et al. 2020; Oksdath Mansilla et al. 2021; Hashmi et al. 2022). For example, Hashmi et al. 2022 improved live imaging of gastruloids using micro-wells but required reducing gastruloid size to ∼ 50 cells, compared to the ∼ 300 cells in the classical protocol optimized for symmetry breaking and axis elongation (Van Den Brink, Baillie-Johnson, et al. 2014). By contrast, our hydrogel embedding approach is compatible with gastruloids of any size or developmental timing, allowing the use of the optimally defined number of seeded cells to study A-P axis formation and produce the full range of cell populations described in the gastruloid system (Bennabi et al. 2024; Fiuza et al. 2024; Van Den Brink, Baillie-Johnson, et al. 2014). This user-friendly system facilitates long-term imaging without altering gastruloid development. Using gastruloids with a low proportion of H2B-iRFP-expressing cells, we successfully tracked individual cell movements without the need for complex image registration (Figure 4B, Figure S7A,B).

**Figure 4:**
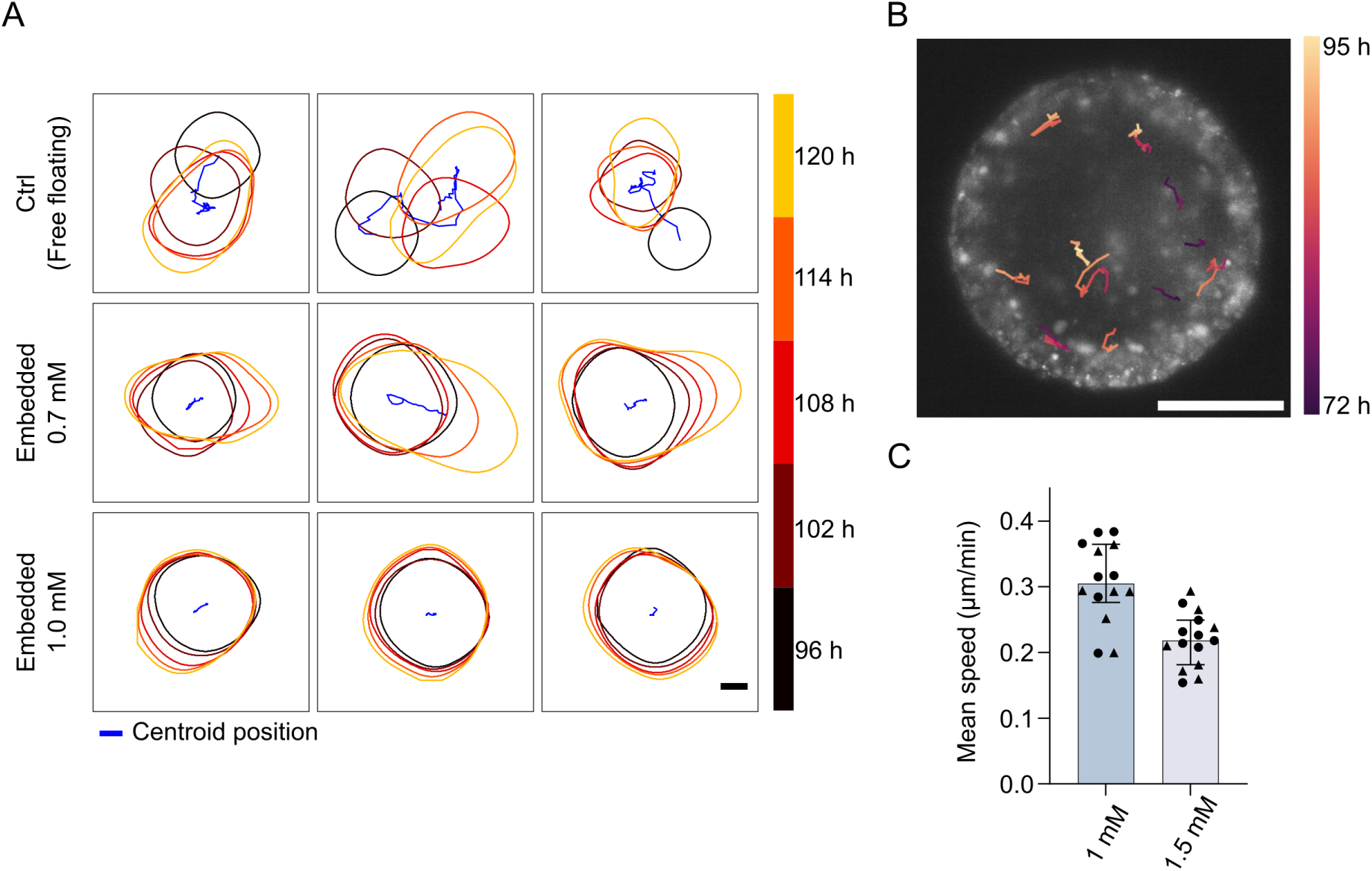
Impaired cell motility in dense gel confinement. (A) Outlines of developing gastruloids from 96 h to 120 h (five time points encoded by color) post seeding under control (freely floating) or embedded (0.7 mM or 1.0 mM) conditions. The trajectory of the gastruloid centroid is shown in blue. Scale bar: 100 μm (B) Colour-coded tracks of cell migration in a gastruloid embedded in a 1.0 mM gel, imaged from 72 h to 95 h post seeding. The image shows H2B-iRFP fluorescence at 95 h post seeding. Scale bar: 100 μm. (C) Mean cell speed for tracks with durations ranging from 3 h 20 min to 8 h 20 min, comparing conditions of 1.0 mM and 1.5 mM gel embedding. Data points represent individual tracks from two gastruloids per condition (triangle and circle symbols). Sample sizes: 1.0 mM (*N* = 14), 1.5 mM (*N* = 15).

To investigate whether impaired cell migration could explain the defects in BRA/SOX2 pole formation observed in 1.5 mM gels, we conducted live imaging and cell tracking in gastruloids embedded at 72 h in either 1.0 mM or 1.5 mM hydrogels. Imaging was performed using the LS2 Viventis system, with adaptations to the gel embedding protocol to accommodate the sample holder (see Methods). Gastruloids were generated using a mixture of cells expressing either a BRA reporter (TProm-mVenus) or a nuclear marker (H2B-iRFP), enabling cell migration tracking and confirmation of BRA expression patterns (see supplementary movies).

Tracking over several hours revealed significantly lower mean cell migration speeds in 1.5 mM gels compared to 1.0 mM gels (Figure 4C). As the organoids grew during the acquisition period, with more growth observed in 1.0 mM gels, we measured radial growth rates to exclude the possibility that differential growth contributed to the observed differences in cell migration speeds. Radial growth was minimal (1.0 mM: 0.015 ± 0.002 μm/min; 1.5 mM: 0.004 ± 0.006 μm/min) and could not account for the substantial reduction in cell migration speed (Figure 4C).

These results suggest that the inability of gastruloids embedded in 1.5 mM gels to establish a BRA/SOX2 pole is not due to transcriptional changes but rather to impaired cell motility and morphogenesis within the mechanically constrained environment. By altering the mechanical properties of the hydrogel, we were able to disrupt polarization and morphogenetic processes while preserving transcriptional profiles, underscoring the critical role of cell migration in BRA/SOX2 pole formation.

## Discussion

This study addresses a key challenge in developmental biology: disentangling the mechanical and chemical contributions of the environment to gastruloid development, an issue often confounded by the variability and undefined composition of traditional matrices such as Matrigel. By employing a bioinert hydrogel system with tunable stiffness and embedding timing, we provide a precise platform for probing how external mechanical constraints influence developmental processes, including elongation, polarization, and transcriptional regulation.

We demonstrated that gastruloids embedded in ultra-soft hydrogels (*<* 1.0 mM) elongate robustly while maintaining AP axis patterning and transcriptional profiles similar to controls. This finding highlights the capability of our system to support nearly unaffected developmental processes in a mechanically controlled environment. Beyond developmental outcomes, the hydrogel’s stability minimizes sample movement during live imaging, enabling precise long-term tracking of gastruloid dynamics. Unlike existing solutions such as micro-wells, which often impose size constraints or require extensive image registration, our hydrogel platform works irrespective of gastruloid size or developmental timing, allowing the use of optimal cell numbers for studying axis formation (Van Den Brink, Baillie-Johnson, et al. 2014; Bennabi et al. 2024; Fiuza et al. 2024). Furthermore, the bioinert nature of our hydrogel enables the disentangling of mechanical and chemical effects, while its modular design allows for functionalization to explore cell-ECM interactions systematically.

A key finding of this study is the decoupling of transcriptional profiles and AP axis patterning under specific mechanical constraints. Embedding in stiffer hydrogels disrupted polarization without altering transcriptional profiles, while earlier embedding significantly affected transcription independently of polarization defects. For example, altering embedding timing impacted the expression of key developmental genes such as *T/Bra*, without disrupting polarized patterning. This suggests that the transcriptional program proceeds autonomously within each cell, relying on local or short-range cues rather than long-range gradients. These findings challenge conventional views of tight coordination between transcription and patterning and raise new questions about how these processes interact under distinct mechanical conditions.

We also identified cell motility as a key factor in polarization, likely mediated through cell sorting and aggregation mechanisms. Gastruloids embedded in stiffer hydrogels (1.5 mM) exhibited significantly reduced cell motility compared to those in softer hydrogels (1.0 mM), potentially explaining their inability to establish a BRA/SOX2 pole. Increased cell density within aggregates in stiffer hydrogels may contribute to reduced motility, but additional factors such as altered cytoskeletal dynamics, signaling pathways, or changes in cell adhesion properties are also likely involved (Cermola et al. 2022; Underhill and Toettcher 2023; Jong et al. 2024; Anlaş et al. 2024; Mayran et al. 2023). These findings underscore the importance of cell migration in axis polarization and highlight the platform’s potential for exploring how mechanical constraints shape cell behavior and morphogenesis.

While our experiments reveal important insights into the interplay of transcription, patterning, and motility, they cannot definitively establish whether gene expression and patterning are entirely independent processes. The observed uncoupling may reflect specific developmental stages or conditions rather than a universal principle. Similarly, the role of motility in polarization warrants further investigation to clarify the causal relationships among these processes. Nevertheless, our hydrogel system provides a robust framework to systematically test these interplays and disentangle the mechanisms underlying early development. By offering precise control over the mechanical environment, this platform opens new avenues for probing the regulatory principles that guide embryogenesis.

## Author contributions

JWN, SG and TG conceptualized the work. JWN, JP, AS, DD, SG, and TG designed experiments and developed experimental protocols. JWN and JP performed experiments and computational image analysis. AM generated the fluorescent reporter cell lines. AM, LD, and PO performed bulk RNA sequencing and analysis. JWN, JP, and TG wrote the manuscript, with input from LLD and AM. DD, SG, and TG secured funding and supervised the work.

## Acknowledgments

We thank I. Bennabi, M. Cerminara, C. Chureau, L. Friedman, M. Lutolf, M. Merle, and P. Hansen. We acknowledge the use of AI-based writing tools for language enhancement. This work was supported by Institut Pasteur (particularly the HPC core facility), Centre National de la Recherche Scientifique, the French National Research Agency (ANR-20-CE12-0028’ChroDynE’ and ANR-23-CE13-0021’GastruCyp’ and ANR-10 LABX-73’Revive’), and by funding from the European Research Council (ERC-2023-SyG, Dynatrans, 101118866). In addition, the Ecole Polytechnique Féedéerale de Lausanne also supported this work, the Swiss National Science Foundation (SNSF, grants 310030-196868, CRSII5-189956, and 407940-206405), and the Human Frontier Science Program (HFSP LT000032/2019-L).

## Data and resource availability

The code used to produce the next-generation sequencing analysis (bulk RNA-seq) can be found on: https://github.com/lldelisle/allRNAseqScriptsFromPineauWongNgEtAl20 The bulk RNA-seq data have been deposited to GEO under the accession GSEXXXX.

## Materials and Methods

### Cell culture

Mouse embryonic stem cells (mESCs) were cultured in 6-well plates (TPP) coated with 0.1 % gelatin in water, in a humidified incubator (5 % CO2, 37°C). Cell culture media was prepared as follows: DMEM 1X + Glutamax (Fisher 11584516) supplemented with 10 % Decomplemented FBS (Gibco, 11573397, decomplemented 30 min at 56°C), 1X Non-essential amino acids (NEAA, Gibco 11140-035), 1 mM Sodium Pyruvate (Gibco, 11360-039), 1 % Penicillin-Streptomycin (Gibco, 15140-122), 100 μM 2-Mercaptoethanol (Gibco 31350-010), 10 ng/mL Leukemia Inhibitory Factor (LIF, Miltenyi Biotec 130-099-895), 3 μM GSK3 inhibitor CHIR 99021 (Sigma, SML1046), 1 μM MEK inhibitor PD 035901 (Sigma, PZ0162). Cells were passaged every other day (detached using Trypsin), and experiments were done using cells that were kept in culture for at least two passages or 5 days after thawing. Cells were tested for mycoplasma contamination using the Eurofins Mycoplasma check on a regular basis.

Unless stated otherwise, experiments were performed using the 129/svev mouse embryonic stem cell line.

### Cell line generation

The other cell lines used (TProm-mVenus and TProm-mVenus/H2B:iRFP) were generated at EPFL. To generate the Tprom-mVenus cell line, a region of 1392bp surrounding the *brachyury* gene was selected to monitor the activity of the *brachyury* promoter (see genome browser view of ATAC-seq and RNA-seq in the supplementary materials). For the transgenic assay, a sequence (see supplementary materials) was inserted by gateway cloning (LR reaction) into the SIF-seq construct (Dickel et al. 2014, Addgene plasmid: 51292: pSKB1-GW-hsp68-Venus-H19).

To form the crRNA:tracrRNA duplex, Alt-R CRISPR-Cas9 crRNA (guide sequence: GTTTTAAGATTTCTTTATGG, ordered from idt) and Alt-R CRISPR-Cas9 tracrRNA (IDT) were hybridized in equimolar concentration in nuclease free IDTE buffer (44 μM) by incubation at 95°C for 5 minutes and allowed to cool at 15-25°C on the bench top. The crRNA:tracrRNA duplex were then used to form the RNP complex with a final concentration of 18 μM of Alt-R Cas9 enzyme (previously diluted in Resuspension buffer R from Neon System kit) and 22 μM of crRNA:tracrRNA duplex. The RNP complex was incubated for 15 minutes at 15-25°C.

Embryonic stem cells (E14tg2a) were thawed and kept for one passage. They were then dissociated and washed twice with PBS. 400 000 cells were resuspended in 22 μL of buffer R (Neon System kit) with 2 μL of RNP complex solution and 4 μg of the SIF-seq construct containing the 1392bp near the *brachyury* promoter indicated above. Cells were electroporated with the Neon electroporation system (with 1100 V, 20 ms and 2 pulses settings) and seeded in a 6 cm petri dish with 3 mL of DMEM media. Media was renewed the next day. 24 h later, medium was replace by DMEM complemented by adding 1x HAT selection supplement (Gibco) media and renewed daily for 5 days. 17 clones were picked and allowed to recover for 1 week in DMEM medium supplemented with HT. They were analysed by PCR (mVenus FWD: caccatggtgagcaagggcgag; mVenus REV: ttctgctggtagtggtcggcga; Ampicillin FW: ctgcaactttatccgcctcc; Ampicillin REV: gtgcacgagtgggttacatc).

Positive clones with presence of mVenus and absence of Ampicillin were amplified and independently verified.

To add H2B:iRFP fluorescent protein, 2 μg of pCAG-H2BtdiRFP-IP (gift from Maria-Elena Torres-Padilla, Addgene plasmid # 47884 ; http://n2t.net/addgene:47884 ; RRID:Addgene 47884) was transfected in 300 000 *brachyury* promoter reporter E14tg2a using FuGENE (Promega). Media was changed the following day and 24 h after, puromycin selection was performed over a period of 5 days. The pool of resistant colonies (*>*1000 colonies) were allowed to grow and was passaged as a pool.

### Gastruloid generation

Gastruloids were generated as described in Beccari et al. 2018. The N2B27 medium was prepared every 3 weeks in-house using 250 mL DMEM/F12+GlutaMax (Gibco, 10565018), 250 mL Neurobasal (Gibco, 21103049), 2.5 mL N2 (Gibco, 17502-048), 5 mL B27 (Gibco, 17504-044), 1X Non-essential amino acids (NEAA, Gibco 11140-035), 1 mM Sodium Pyruvate (Gibco, 11360-039), 100 μM 2-Mercaptoethanol (Gibco 31350-010), 1 % Penicillin-Streptomycin (Gibco, 15140-122), 2.5 mL Glutamax (Gibco, 35050061). Gastruloids were generated by manually seeding 300 cells per well in Costar Low Binding 96-well plates (Costar, Corning, 7007), in a volume of 40 μL per well. After 48 h of aggregation, spheroids were exposed to a 24 h pulse of Wnt agonist by adding 150 μL of 3 μM CHIR 99021 (CHI in the text) in N2B27 to each well, unless stated otherwise. N2B27 media was then changed every 24 h by replacing 150 μL of media per well until 120 h. For gastruloids in hydrogels, media was also replaced every 24 h.

### Gel embedding

Gastruloids were embedded in hydrogels 72 h or 96 h after seeding, then left to grow until 120 h. For gastruloid embedding, 20-30 gastruloids were collected from a 96-well plate and left to sediment in a falcon tube. Meanwhile, gel components were prepared, following the proportions of the manufacturer of the 3-D Life Dextran-PEG Hydrogel FG (Cellendes, FG90-1, containing the components PEG-Link, Dextran-Maleimide and CB Buffer pH=5.5) to reach the indicated function concentrations, setting 50 % of gastruloid suspension and a total volume of 120 μL per gel. For each gel, two tubes were prepared: one tube containing ultrapure water and PEG-Link, and one tube containing CB Buffer (pH=5.5) and gastruloid suspension in N2B27. Dextran-Maleimide was added to the second tube at the last moment, followed by a quick homogenization with the pipette, pooling of the two solutions, homogenization, then deposition of the gel mix into a glass-bottom dish (Cellvis, D35-10-1.5-N). Quickly, a membrane was deposited on top of the gel (Isopore filter, 5.0 μm membrane, 13 mm diameter, Merck, TMTP01300), followed by an adhesive ring (Delta microscopies, slide wells D70366-12) to secure the membrane. The dish was then put in the incubator for 15 min for gel formation, after which 2 mL of N2B27 was added on top, and the embedded gastruloids were kept for culture as usual.

### Gel characterisation

Hydrogel stiffness was measured using a rheometer (Kinexus Ultra, Malvern). Briefly, gel formation was measured by preparing the gel mix as described above, only replacing the gastruloid suspension by N2B27 media. Upon mixing, 100 μL of gel mix was deposited between the rheometer geometry (flat 20 mm diameter tool) and plate, kept at 4°C. Excess of liquid was removed. Temperature was quickly ramped up to 37°C while oscillating rotations of the mobile geometry measured the gel stiffness. Frequency and amplitude were fixed at 3 Hz and 5 %. After 15 min, N2B27 was added in contact of the gel to simulate the effect of adding medium in the usual experiment. Gelling dynamics could be observed through the shear modulus measurement in live, as shown in Supplementary Figure 1. A stable plateau value was reached within 30 minutes.

### Bulk RNAseq sample preparation and Analysis

#### Sample generation

Gastruloids were generated and embedded as described above, and as a negative control, samples where the mESCs aggregates were not submitted to a CHI pulse and left in a 96-well plate were generated. For each replicate, about 30 gastruloids were processed per condition. At 120 h after seeding, dishes containing embedded gastruloids were treated as follow: the adhesive ring was lifted, and a solution of 1:20 Dextranase (Cellendes, D10-1) in PBS (Ca^++^/Mg^++^) was added to each dish, and left *>*20 min in the incubator until gastruloids were freely moving and could be collected in a falcon tube. Gastruloids in 96-well plates (control and negative control) were collected into falcon tubes. All gastruloids were then washed twice with cold PBS (Ca^++^/Mg^++^) before being snap-freezed in liquid nitrogen, and stored at −80°C until RNA extraction.

#### Sample processing

RNeasy Mini kit (Qiagen) with on-column DNase digestion was used for RNA extraction following manufacturer’s instructions. RNA quality was assessed on a TapeStation TS4200, all RNA samples showed quality number (RIN) above 9. RNA-seq library preparation with Poly-A selection was performed with 550 ng of RNA using the Illumina stranded mRNA ligation and following the manufacturer’s protocol 1000000124518 v01. Libraries were quantified by qubit DNA HS and profile analysis was done on TapeStation TS4200. Libraries were sequenced on Novaseq 6000, with paired end 75 bp reads.

#### bulk RNAseq analysis

RNA-seq preprocessing was done using a local installation of Galaxy (The Galaxy Community 2024). Adapter and bad quality bases were removed from fastq files using cutadapt version 4.4 (Martin 2011) (-q 30 -m 15 -a CTGTCTCTTATACA-CATCTCCGAGCCCACGAGAC -A CTGTCTCTTATACACATCTGACGCTGC-CGACGA). Filtered reads were aligned on mm10 using STAR version 2.7.10b (Dobin et al. 2013) with the ENCODE parameters and a custom gtf (Lopez-Delisle 2021). FPKM were computed with cufflinks version 2.2.1.3 (Trapnell et al. 2010; Roberts et al. 2011) using –max-bundle-length 10000000 –multi-read-correct –library-type ”fr-firststrand” -b mm10.fa –no-effective-length-correction -M mm10 chrM.gtf. For analyses, genes from mitochondrial genes were excluded. For both analyses (concentration effect and time effect), the FPKM values were transformed with log2(1+FPKM) and the 500 genes with the highest variance were selected. PCA was computed on these genes and clustering was performed using 1 - Pearson’s correlation coefficient as distances with ward.D2 method. Pairwise differential expression analysis was computed with DESeq2 (Love, Huber, and Anders 2014) on raw counts from STAR (excluding mitochondrial genes). A gene was considered as differentially expressed when the adjusted p-value was below 0.05 and absolute log2 fold-change above 1.

### Transmitted light imaging for morphological characterisation

At 120 h after gastruloid seeding, embedded gastruloids or gastruloids cultured in 96-well plates were imaged in transmitted light using an Olympus video-microscope with the Olympus CellSens dimension 3.1 software, equipped with a Hamamatsu C11440-36U CCD camera with a pixel size of 5.86*5.86 μm and a 4X 0.13 NA objective. Gastruloids in 96-well plates were imaged individually, whereas gastruloids in gels were imaged by tiling over the whole region of the glass-bottom dish, then performing stitching on Fiji (Schindelin et al. 2012). Embedded gastruloids could then be cropped from the large region to obtain individual TIFF images of embedded gastruloids.

Images were then processed in Python to obtain a mask of each gastruloid and measure morphological characteristics such as the gastruloid length (obtained by computing the medial axis and extending it to the organoid extremities, as described in Merle et al. 2023, the straightness ratio (ratio between the gastruloid length and the distance between the body axis extremities), and the aspect ratio.

### Gastruloid immunostaining

For this protocol, only PBS (Ca^++^/Mg^++^) was used, and will be called PBS in this section. Any tube, plate or pipette tip that contained gastruloids was either low-binding or coated with PBSF (PBS, 10 % FBS). Washes were made by spinning the gastruloids for 1 min at 10 g to help sedimentation, and aspirating the liquid. Gastruloids cultured in a 96-well plate until 120 h were collected in a low-binding 15 mL falcon tube and washed with PBS, before proceeding with 2 h fixation in 4 % PFA in PBS at 4°C. Meanwhile, embedded gastruloids were fixed as follows: N2B27 was removed from the dishes, and gels were washed with PBS before 2 h 30 fixation in 4 % PFA in PBS at room temperature (RT). After fixation, samples were washed twice for 15 min in PBSF (PBS, 10 % FBS) at RT, and once in PBS.

Embedded gastruloids were de-embedded after this step: the adhesive ring and membrane were lifted, and dishes were incubated with 1:20 Dextranase (Cellendes, D10-1) in PBS at 37°C for *>*20 min, until gastruloids could freely move in the dish. Gastruloids were then recovered in a low-binding falcon tube, and washed with PBS.

All gastruloids were then permeabilized by incubating twice for 30 min in 13 mL of PBSFT (PBS, 10 % FBS, 0.03 % Triton X-100) at RT. Primary antibody staining was performed O/N at 4°C by incubating in a solution of 1:200 Rat anti SOX2 (eBioscience, 14-9811-80), 1:200 Rabbit anti BRACHYURY (Abcam, ab209665), 1:500 DAPI (Sigma, D9542-5MG), in 500 μL PBSFT (per condition). The next day, gastruloids were washed twice for 20 min at RT in PBSF, once in PBS, then in PBSFT. They were then incubated O/N at 4°C with secondary antibodies, in 500 μL of mix containing 1:500 DAPI, 1:500 Donkey anti-Rabbit AF647 (Invitrogen, A-31573), 1:500 Donkey anti-Rat AF488 (Invitrogen, A-21208) in PBSFT. Finally, gastruloids were washed twice 20 min at RT in PBSF, then in PBS. They were then transferred to a 6-well plate filled with PBS using a cut P1000 tip, to finish washing the gastruloids and remove debris or impurities. Subsequently, gastruloids were transferred to 1.5 mL eppendorfs, where all PBS was removed before resuspending in 150 μL of mounting media (50/50 PBS/Aquapolymount, Polysciences 18606-20), and transferring them to a glass-bottom dish (Cellvis, D35-10-1.5N). Gastruloids could then be moved to scatter them across the dish, and were left to sediment at the bottom of the dish for *>*24 h at 4°C, after sealing dishes with Parafilm to minimize evaporation. Samples were then sealed with a coverglass and nail polish for conservation.

### Confocal imaging of immunofluorescence samples

Immunofluorescence samples were imaged using a Zeiss LSM980 confocal microscope controlled with the Zen 3.3 software (Zeiss), and equipped with a 10x 0.45 NA air objective (Zeiss). Images were acquired as z-stacks, by taking 30 slices in a 150 μm range, resulting in a voxel size of 0.22 x 0.22 x 5.00 μm.

### Extraction of 1D gene expression profile

Intensity profiles along the antero-posterior axis were computed as described in Merle et al. 2023. All analysis was performed on maximum intensity projections of confocal images stacks. Using a custom python script, masks of gastruloids were generated from the DAPI channel. The main body axis of the gastruloid was then defined by finding the medial axis and expanding its extremities with straight lines, tangent to the medial axis ends, until the lines cross the gastruloid contour. The contour was cut at these intersection points, and each side was subdivided in 100 equidistant points. Sections of the gastruloids could be defined by connecting pairs of equivalent points on each side. To obtain the intensity profiles of fluorescent signals, the average intensity in each bin was measured. Since the fixation protocol is different for embedded and non-embedded gastruloids, profiles were compared by shape, and not fluorescence intensity, and were therefore normalized as follows. For each profile, the average of the 10 % lowest values and the average of the 10 % highest values were taken, and the profile was normalized to these values.

### Live imaging of gastruloid elongation and pattern formation by videomicroscopy

Live imaging of BRACHYURY pattern dynamics and gastruloid elongation with the TProm-mVenus cell line were performed with a wide-field inverted fluorescence microscope (IX81, Evident) using a 20x objective (UPLFLN20X). Gastruloids were embedded in 0.7 μM dextran-gels at 96 h. For each embedded gastruloid, z-stacks with 20 μm spacing were acquired every 30 minutes in bright field and fluorescence. Gastruloid contour and medial axis were extracted from brightfield images and intensity profiles were extracted along this axis.

### Cell tracking

Movies for single cell tracking were acquired on gastruloid chimeras composed of TProm-mVenus and TProm-mVenus/H2B:iRFP. Images were acquired every 20 min starting from 72 h, using a LS2 Viventis light sheet microscope with its 25x objective configuration. This resulted in a voxel size of 0.26×0.26×3 μm. Cell tracking was performed by manual tracking using Trackmate on the Fiji software in 3D (Schindelin et al. 2012). Gels were cast in the LS2 viventis sample holders (SHT SW0.8) without a membrane deposited on top. Movies from 2 gastruloids embedded in a 1.0 mM gel and 2 gastruloids embedded in a 1.5 mM gel were analyzed, with only trajectories of 10 to 25 time points being considered.

### Sequence used in transgenic assay

A region of 1392bp surrounding the brachyury was selected to monitor the activity of the brachyury promoter (see genome browser view of ATAC-seq and RNA-seq below).

ACCCCAGAGGTTGGCTCCTGGAAAACCGTCTCTCCCAGAAGTAGGGGCAGGTAGAACCCACAACTCCGACCCCAAAGACTTCCCAGGGAGACTCTCAGAGAGACAACGAACTCAGAATTGAGTGCCCCCACCTGATTCAGGGGCCTCTTCCAAGGAGCTTCGGGATAGGATAGGAGAGTGGAAGACGGGGAAACGGAGGCTGGAACCCAGAGTCTCGTTAAAGAGCTGGGCGCGAGCTCTGGCTTCCTTCCCGCTTCCTGGGCTCCCGTTTTAGAGGAATGTTATTGTTTAAAGAGACCCCATTGAACTATTTCCTGCTCTTTGTCACCTTCCCCTCACTCTCCCGGCAGAGGTTCTCACCGAGAGGCAATAAACCAACTGCTGCCCACACCGCATGGCGAGGCGGGTAGGGAAACGCGCGCAGCATGCGTTCCAACAATCCCCGGCGCAAAGAGACCAGGGACTCCCGGGGCCACATTCGGTGCAGGCGCATCCACCGTCAAAGTCCAGCTTTTATGTGGGACGCGAGGACACCTCCTACTAGGGTCGCTATCTGTTCGTCTATTTCCCTCTCTGGACAGATCCGCATTGAGCTTCCCTCTCCACGCAGGTGAAGGTCGTGGGGGACCTGGATGCCGAGGTGGGAGTTAGTGGCAGTCCATGGGGCGAGGGGACGTGTCCCAAAGCTGCCACACCTGGGGAGGCTGAGGCTTTGGAGAGGTCAAGGAGACCCGGGAGACGCCGATCCGCCGAAGTCCCTCTCAGGTGCGCGCAGCGTGGACACTCCGCGGGGCAAAGTCGCAGGCGCCGGTGTGCGCTTGGACAGCGCGTGGGAGTGGAGAGTTTAGCAGTGGCTCTAGGAGCCAGGGTCCTGGGTGGCTCCAGCCCGGCTTCTCGCCCTCCCTCCCCCAGGGTCCGCCCCGCCGCTTTGATGGAGGTGCAAACATTTGGGGGAGGGCGGGGGTGTCGGGACTGCGCCCGACGCTTTCCTTACAGGAAGCGCGCGCTGGAGCCCATTGTTGGCCCCCAGCCTCCGGGCCCGCCCGGCCAGTCTGATATGGCCGCGCACCGCCAATGGGCAGCTGCTCGGTACTTCAAAGGGTGTCCCGCCCAATCCGCCGCACCCCCCTGCGAGGCCACCTCGGCTGTATTTATGGGGAGGGGACCCATTTTTCTCTTCCCCAGAGACTTACTCTTGTCGCGCCTTGCGGGAGTTCAAGTGGAGCCACGGCTCCCCAGGCCCTCTCCCCCATCCCCGCCCCCTTCCCCCCTCATCCCGATCTCGGTGCTCCTTTGGCGAATGTGCAGGGACCCAGGTGTAATCTTTGGGCTCCGCAGAGTGACCCTTTTTCTTGGAAAAGCGGTGGCGAGAGAAGTGAAGGTGGCTGTTGG

### Genome browser view of the brachyury promoter

Bigwig files with RNA-seq profiles of controls gastruloids of 48 h to 120 h were retrieved directly from GEO (GSE247508). Bigwig files with ATAC-seq profiles of wild-type gastruloids of 48 h to 120 h were retrieved directly from GEO (GSE247507). The plot (Figure S8) was generated with pyGenomeTracks (Lopez-Delisle et al. 2020) version 3.9 on mm10:chr17:8,428,652-8,442,571.

---

**Figure S1:**
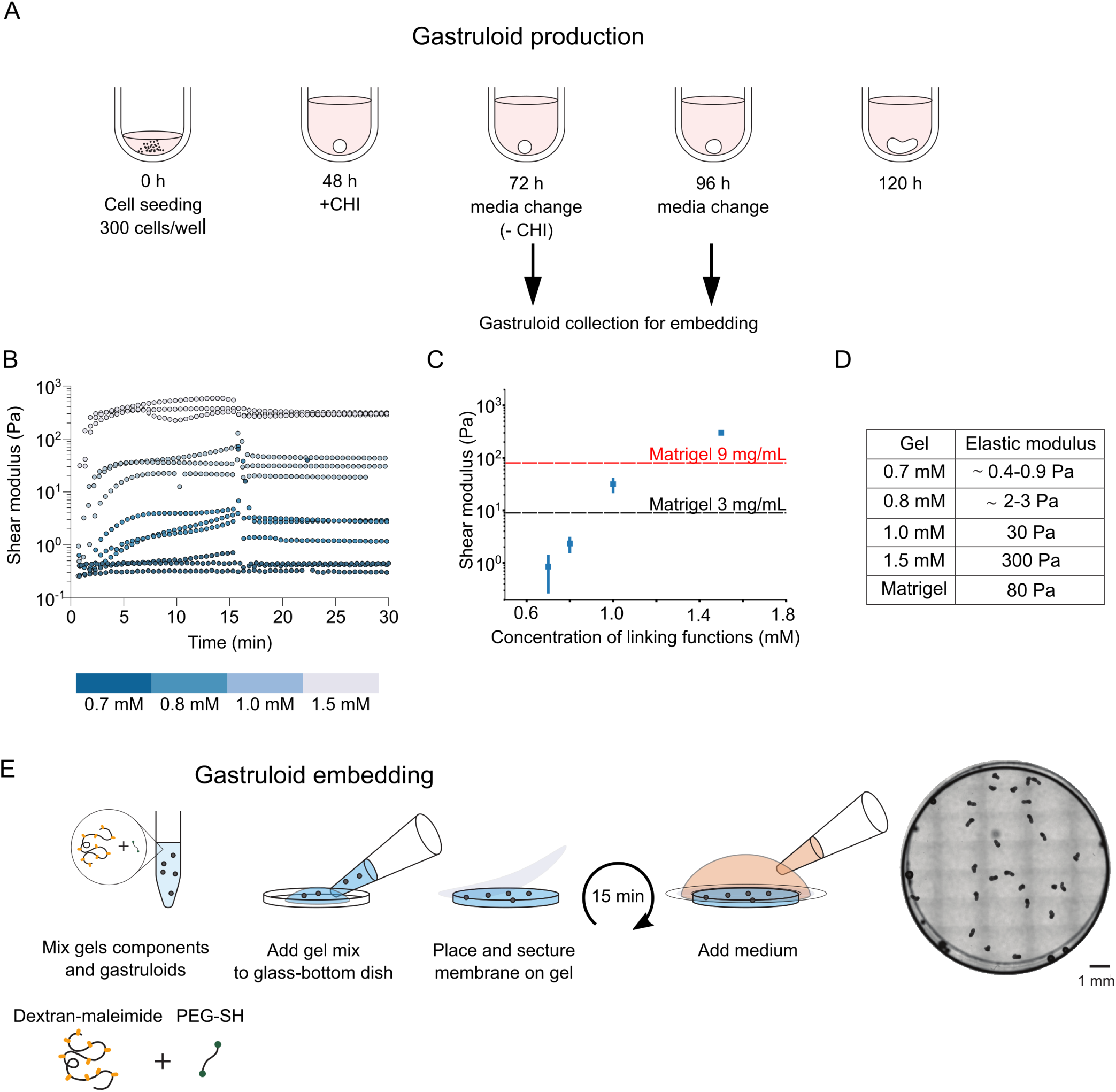
Embedding gastruloids in an ultra-low stiffness dextran-based hydrogel. (A) Schematic view of the gastruloid generation protocol. (B) Gelation dynamics of hydrogels prepared with different concentrations of components (concentration of reactive functions). (C) Stiffness of hydrogels prepared with different concentrations of components (concentration being the concentration of reactive functions), measured using a rheometer. (D) Table with elastic moduli of hydrogels as a function of concentrations of reactive functions. (E) Left: Protocol of gastruloid embedding, using a mix of Dextran-maleimide and PEG-SH. Right: Overview of a dish of embedded gastruloids, at 120 h.

**Figure S2:**
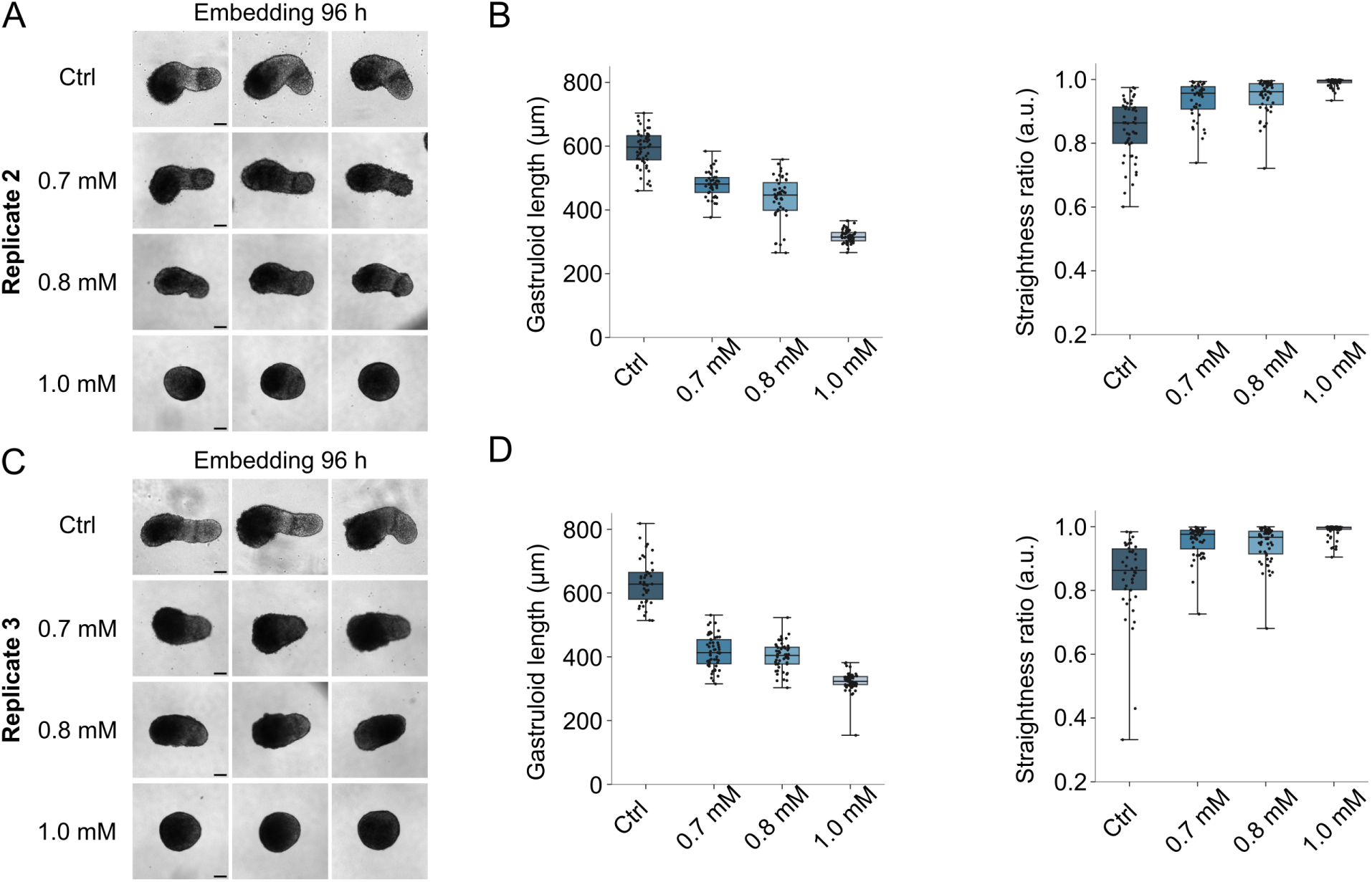
(A) Replicate 2: Bright field images of gastruloids 120 h after seeding, either grown in 96-well plates or embedded in hydrogel at 96 h after seeding. Gel concentrations 0.7 mM, 0.8 mM or 1.0 mM. Scale bar 100 μm. (B) Replicate 2: Length and straightness of gastruloids 120 h after seeding, for gastruloids either grown in 96-well plates or embedded in hydrogel at 96 h after seeding. Gel concentrations 0.7 mM, 0.8 mM or 1.0 mM. Data obtained from bright field images as represented in A: Ctrl N=52; 0.7 mM N=42; 0.8 mM N=48; 1.0 mM N=59. (C) Replicate 3: Bright field images of gastruloids 120 h after seeding, either grown in 96-well plates or embedded in hydrogel at 96 h after seeding. Gel concentrations 0.7 mM, 0.8 mM or 1.0 mM. Scale bar 100 μm. (D) Replicate 3: Length of gastruloids 120 h after seeding, for gastruloids either grown in 96-well plates or embedded in hydrogel at 96 h after seeding. Gel concentrations 0.7 mM, 0.8 mM or 1.0 mM. Data obtained from bright field images as represented in C: Ctrl N=42; 0.7 mM N=56; 0.8 mM N=52; 1.0 mM N=68.

**Figure S3:**
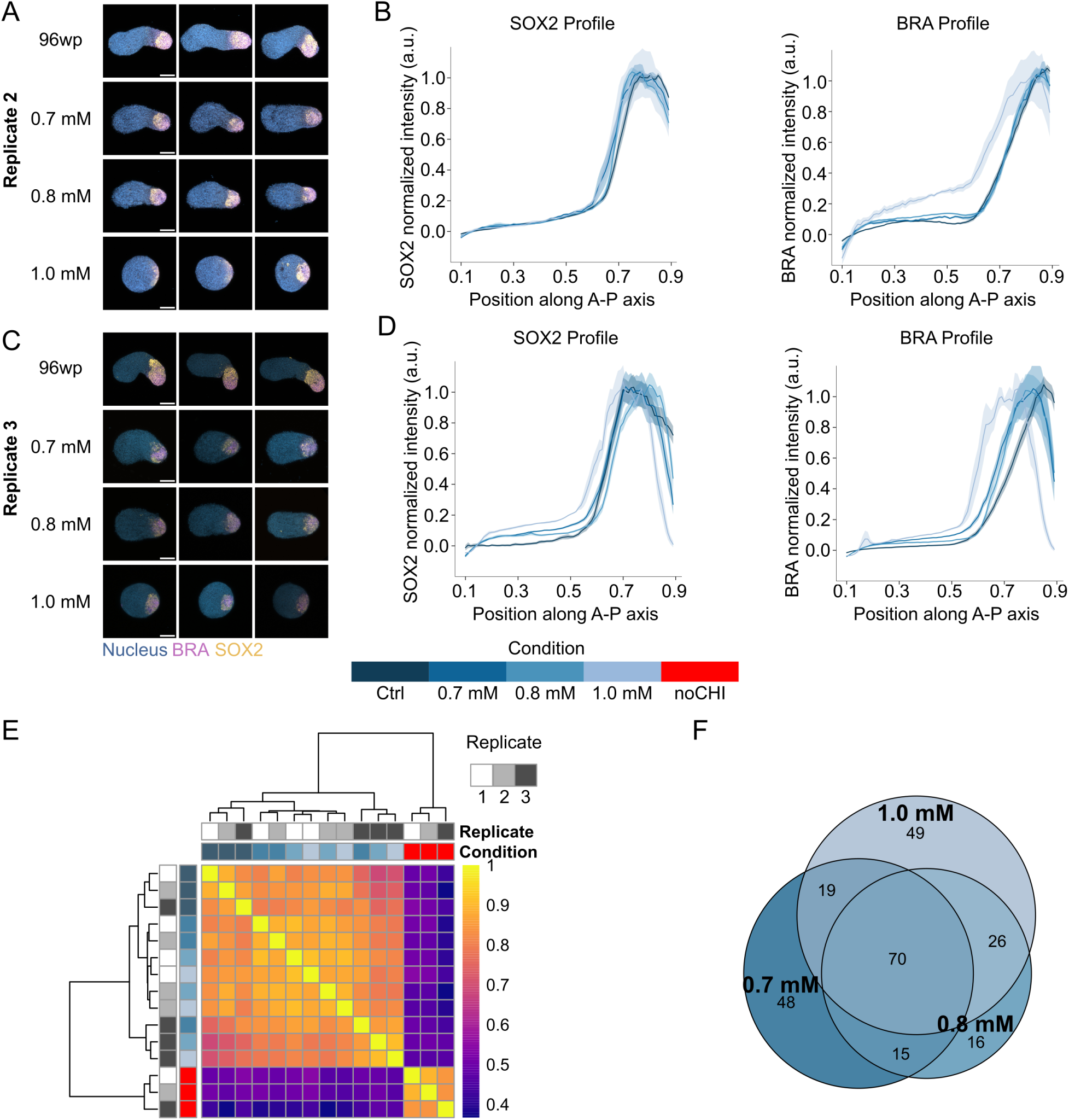
(A) Replicate 2: Immunofluorescence images of gastruloids 120 h after seeding, either grown in 96-well plates or embedded in hydrogel at 96 h after seeding. Gel concentrations 0.7 mM, 0.8 mM or 1.0 mM. Blue: Nucleus, Purple: BRA, Orange: SOX2. Scale bar 100 μm. (B) Replicate 2: Normalized expression profiles (Mean ± SEM) of SOX2 and BRA along the AP axis. Ctrl N=26; 0.7 mM N=14; 0.8 mM N=20; 1.0 mM N=8. (C) Replicate 3: Immunofluorescence images of gastruloids 120 h after seeding, either grown in 96-well plates or embedded in hydrogel at 96 h after seeding. Gel concentrations 0.7 mM, 0.8 mM or 1.0 mM. Blue: Nucleus, Purple: BRA, Orange: SOX2. Scale bar 100 μm. (D) Replicate 3: Normalized expression profiles (Mean ± SEM) of SOX2 and BRA along the AP axis. Ctrl N=21; 0.7 mM N=12; 0.8 mM N=8; 1.0 mM N=3. (E) Correlation matrix of bulk RNA sequencing experiments clustered using the ward-D2 algorithm, where noCHI is used as a negative control for gastruloid formation. Colors indicate the Pearson’s correlation coefficient values. All samples are analyzed in three independent replicates. (F) Euler diagram showing the overlap between DE genes using non-embedded as the control condition.

**Figure S4:**
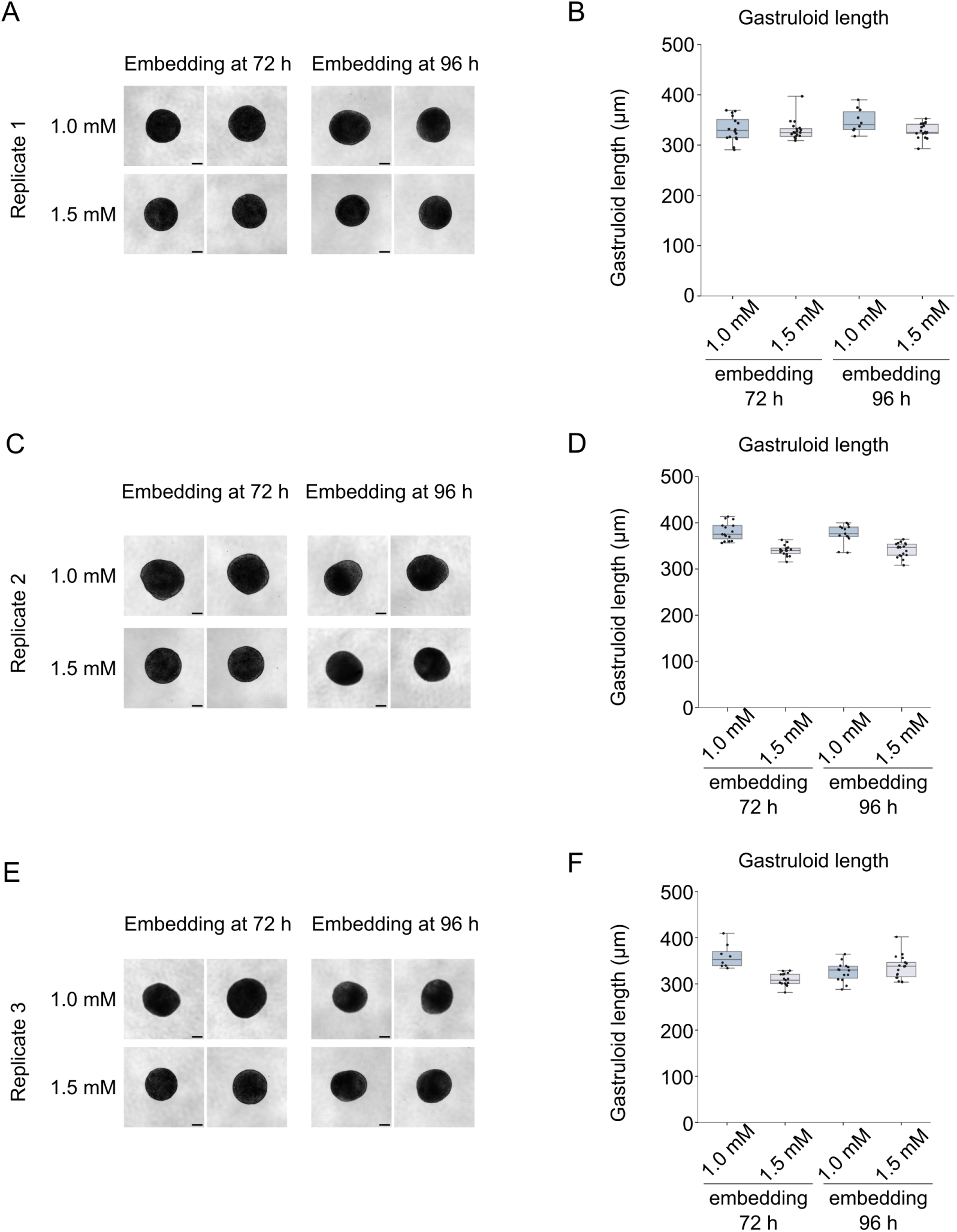
(A) Replicate 1: Bright field images of gastruloids 120 h after seeding, embedded in hydrogel at 72 h or 96 h after seeding. Gel concentrations 1.0 mM or 1.5 mM. Scale bar 100 μm. (B) Length of gastruloids 120 h after seeding, for gastruloids embedded in hydrogel at 72 h or 96 h after seeding. Gel concentrations 1.0 mM or 1.5 mM. Data obtained from bright field images as represented in A, a representative experiment. Embedded 72 h: 1.0 mM N=16; 1.5 mM N=20. Embedded 96 h: 1.0 mM N=10; 1.5 mM N=18. (C) Replicate 2: Bright field images of gastruloids 120 h after seeding, embedded in hydrogel at 72 h/96 h after seeding. Scale bar 100 μm. (D) Replicate 2: Length of gastruloids 120 h after seeding, for gastruloids embedded in hydrogel at 72 h/96 h after seeding. Data obtained from bright field images as represented in C. 72 h 1.0 mM N=15; 72 h 1.5 mM N=16; 96 h 1.0 mM N=13; 96 h 1.5 mM N=16. (E) Replicate 3: Bright field images of gastruloids 120 h after seeding, embedded in hydrogel at 72 h/96 h after seeding. Scale bar 100 μm. (F) Replicate 3: Length of gastruloids 120 h after seeding, for gastruloids embedded in hydrogel at 72 h/96 h after seeding. Data obtained from bright field images as represented in E. 72 h 1.0 mM N=8; 72 h 1.5 mM N=17; 96 h 1.0 mM N=14; 96 h 1.5 mM N=18.

**Figure S5:**
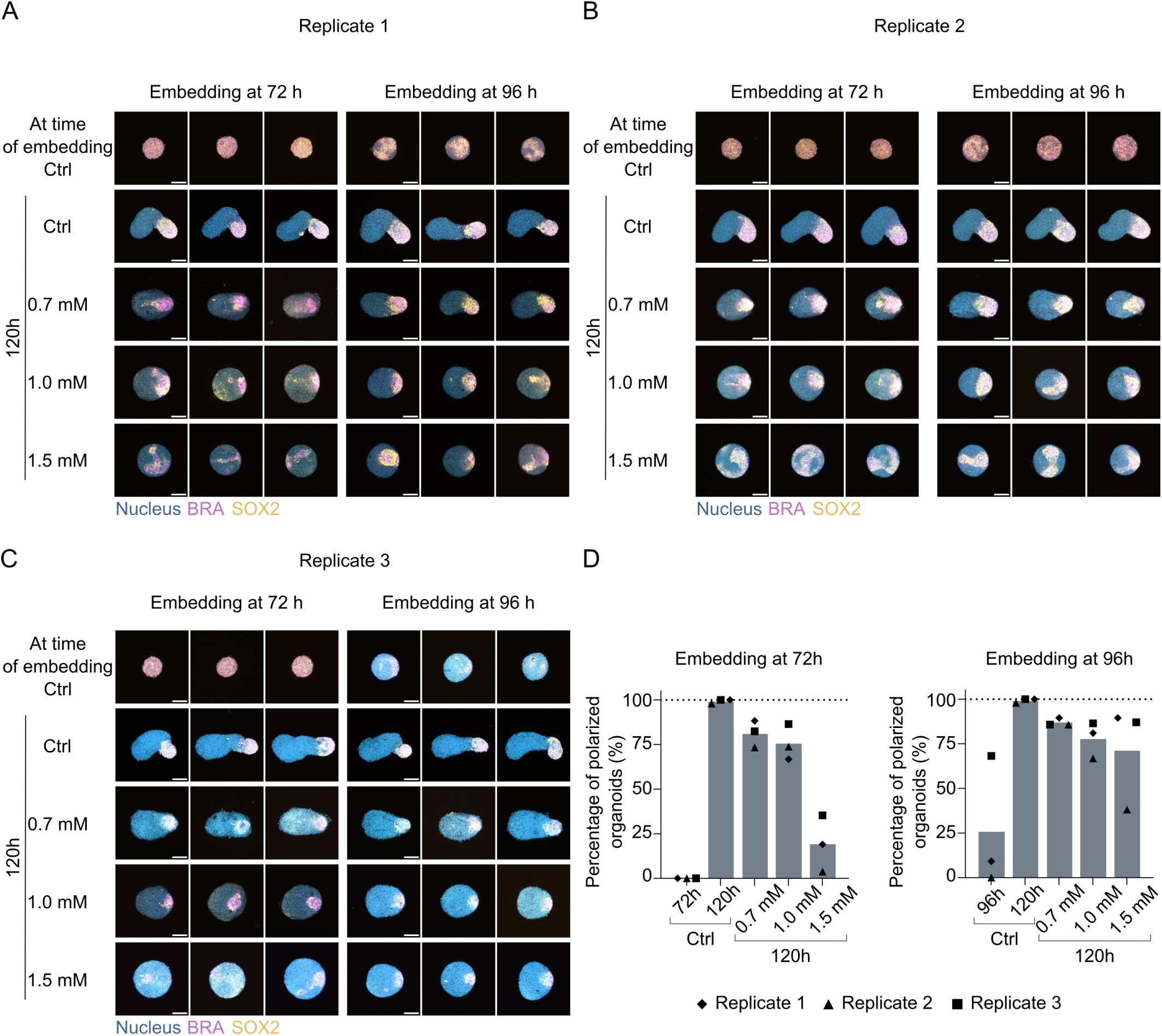
(A),(B),(C) For Replicates 1, 2 and 3 respectively: Immunofluorescence images of gastruloids 120 h after seeding, either grown in 96-well plates (Ctrl) or embedded in hydrogel at 72 h/96 h after seeding. Blue: Nucleus, Magenta: BRA, Yellow: SOX2. Scale bar 100 μm. (D) Quantification of the percentage of gastruloids that formed a unique BRA/SOX2 pole at 120 h or at time of embedding, analyzed from immunofluorescence images as in (A), (B) and (C), for gastruloids embedded at 72 h or 96 h. Number of analyzed gastruloids (per replicate): (Left) 72 h N=20/18/22 ; 120h Ctrl N=25/48/24; 0.7 mM N=17/15/17 ; 1.0 mM N=15/19/22 ; 1.5 mM N=21/27/17.(Right) 96 h N=22/23/22 ; 120h Ctrl N= 25/48/24 ; 0.7 mM N=19/21/21 ; 1.0 mM N=21/21/22 ; 1.5 mM N=19/21/23.

**Figure S6:**
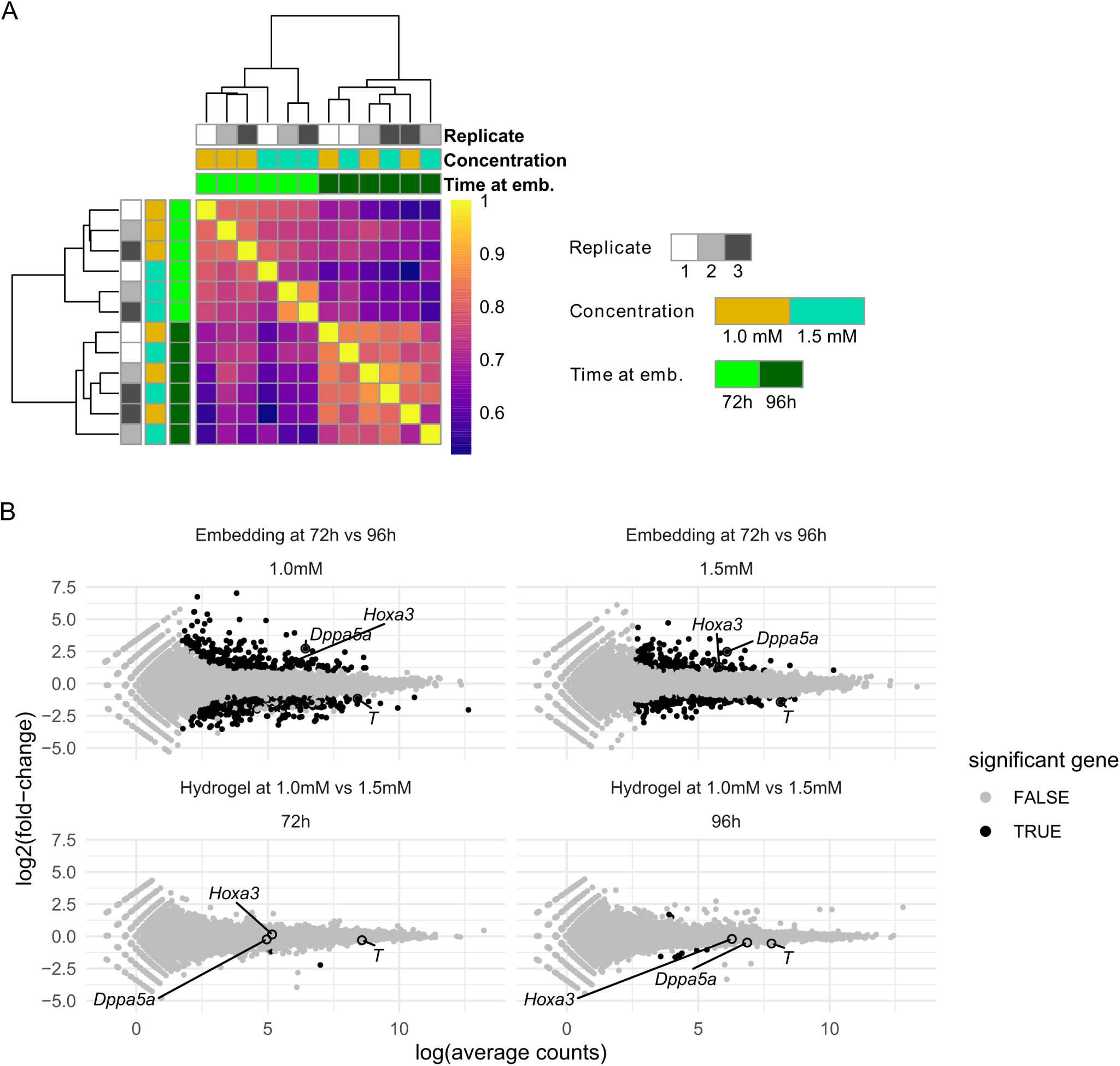
(A) Correlation matrix of bulk RNA sequencing experiments clustered using the ward-D2 algorithm, using samples embedded at 72 h or 96 h, at gel concentrations of 1.0 mM or 1.5 mM. Colors indicate the Pearson’s correlation coefficient values. All samples are analyzed in three independent replicates. (B) MA-plot showing the differential gene expression (log2FC ≥ ± 1, adjusted p-value*<*0.05 as measure by DESeq2 analysis) determined from bulk RNA sequencing. On the upper row, comparing gastruloids embedded at 72 h vs 96 h, for embedding in 1.0 mM or 1.5 mM gels. On the lower row, comparing gastruloids embedded in 1.0 mM vs 1.5 mM gels, for embedding at 72 h or 96 h.

**Figure S7:**
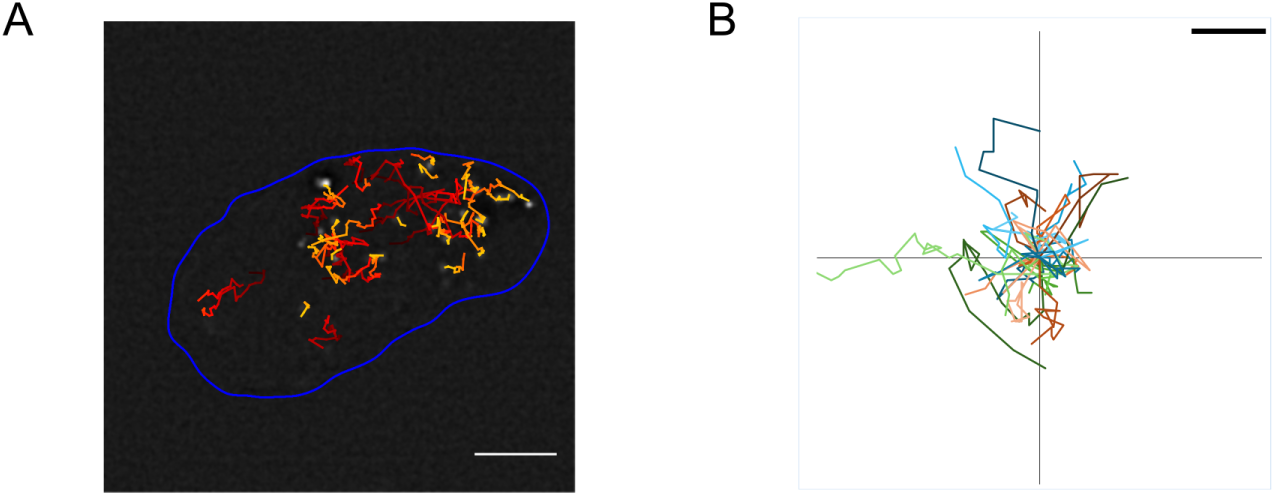
(A) Tracks of cells in a gastruloid embedded in a 0.7 mM gel, filmed from 98 h to 120 h using a classical epifluorescence microscope. Contour in blue was determined using thresholded brightfield image. Tracks were obtained using trackmate. Scale bar 50 μm. (B) Example trajectories of cells tracked in gastruloids embedded in a 1.0 mM gel from 72 h to 95 h. Gastruloids made from a mix of cells expressing either a reporter of BRA expression (TProm-mVenus cells) alone, or together with a nuclear marker (H2B-iRFP). Imaging was done using using the LS2 Viventis system. Scale bar 5 μm.

**Figure S8:**
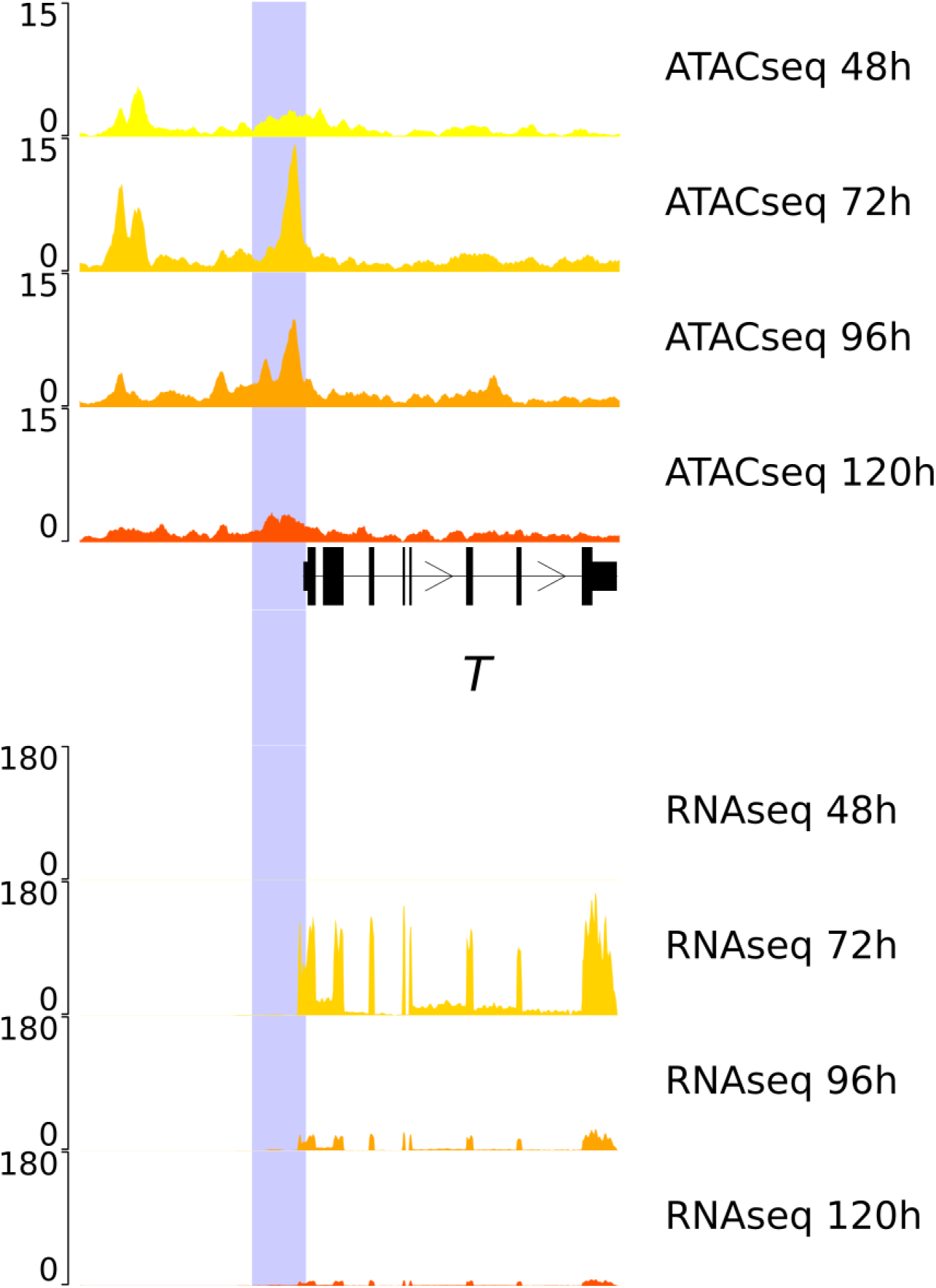
Genome browser view of ATAC-seq and RNA-seq from Mayran et al. 2023 for generation of TProm-mVenus and TProm-Venus/H2B:iRFP cell lines. A region of 1392bp surrounding the brachyury was selected to monitor the activity of the brachyury promoter.

Supplementary table 1: List of differentially expressed genes in the compared conditions the non-embedded condition was used as reference.

Supplementary table 2: List of differentially expressed genes in all pairwise comparisons relative to the time of embedding or to the gel concentration.

